# Estimating the Relative Probability of Direct Transmission between Infectious Disease Patients

**DOI:** 10.1101/612945

**Authors:** Sarah V Leavitt, Robyn S Lee, Paola Sebastiani, Charles R. Horsburgh, Helen E Jenkins, Laura F White

## Abstract

**Background:** Estimating infectious disease parameters such as the serial interval (time between symptom onset in primary and secondary cases) and reproductive number (average number of secondary cases produced by a primary case) are important to understand infectious disease dynamics. Many estimation methods require linking cases by direct transmission, a difficult task for most diseases.

**Methods:** Using a subset of cases with detailed genetic or contact investigation data to develop a training set of probable transmission events, we build a model to estimate the relative transmission probability for all case-pairs from demographic, spatial and clinical data. Our method is based on naive Bayes, a machine learning classification algorithm which uses the observed frequencies in the training dataset to estimate the probability that a pair is linked given a set of covariates.

**Results:** In simulations we find that the probabilities estimated using genetic distance between cases to define training transmission events are able to distinguish between truly linked and unlinked pairs with high accuracy (area under the receiver operating curve value of 95%). Additionally only a subset of the cases, 10-50% depending on sample size, need to have detailed genetic data for our method to perform well. We show how these probabilities can be used to estimate the average effective reproductive number and apply our method to a tuberculosis outbreak in Hamburg, Germany.

**Conclusions:** Our method is a novel way to infer transmission dynamics in any dataset when only a subset of cases has rich contact investigation and/or genetic data.

**KEY MESSAGES:**

- This method provides a way to calculate the relative probability that two infectious disease patients are connected by direct transmission using clinical, demographic, geographic, and genetic characteristics.
- We use a naïve Bayes, a machine learning technique to estimate these probabilities using a training set of probable links defined by contact investigation or pathogen WGS data on a subset of cases.
- These probabilities can be used to explore possible transmission chains, rule out transmission events, and estimate the reproductive number.

## INTRODUCTION

Infectious disease parameters such as the serial interval (time between symptom onset from primary to secondary case) and the reproductive number (average number of secondary cases produced by a primary case over the infection course) are instrumental in managing outbreaks (1). For diseases in which disease progression shortly follows infection, these parameters have been studied extensively (1–6). For others, such as tuberculosis (TB), the serial interval and reproductive number estimates are few and inconsistent (7–9).

Methods to estimate both the serial interval and reproductive number often rely on determining which cases are linked by direct transmission. Pathogen whole genome sequence (WGS) data is a powerful tool to link cases and several methods have been developed to analyze these data (10–21). However, WGS data is still relatively expensive and requires bioinformatics expertise, making universal use in high disease burden settings unfeasible and many datasets are likely to have WGS data on only a proportion of cases. Another way to link cases is contact investigations, which are often part of the infectious disease outbreak response (22–28). However, these often do not perfectly identify due to nonspecific transmission mechanisms, disease characteristics, and the willingness and ability of cases to share information about contacts (25,28–32). In addition, contact investigations are time consuming and require significant human resources, again meaning that this data source is unlikely to be available for all cases.

Here, we present a novel method to predict the relative probability of direct transmission between infectious disease patients using pathogen WGS data and/or contact investigations when these data are only available on a proportion of cases, paired with other risk factor data. These probabilities can be used to understand outbreak transmission dynamics and estimate the reproductive number without a reliable serial interval estimate. We apply our method to a TB outbreak in Hamburg, Germany.

## METHODS

### Data Structure

Our method requires individual-level case data, e.g. geographic location, clinical information, demographics, and sampling date. At least a subset of the cases needs additional information, e.g. detailed contact investigation and/or pathogen genome WGS data, to form the training set to generate the model. We transform this dataset of individuals into a dataset of ordered case-pairs (*i, j*), where case *i* was observed before case *j*. We convert the individual-level covariates (X_1_, X_2_, …, X_p_) into pair-level covariates (*Z*_1_, *Z*_2_, …, *Z*_*p*_) by computing “distances” which capture how well the individuals match on covariates values. For example, if the individual-level covariate *X*_1_ was town of residence, the pair-level covariate *Z*_1_ could indicate if the individuals live in the same town, neighboring towns, or more distant towns. This process is described further in the supplementary material.

### Naïve Bayes

To estimate the probability that cases *i* and *j* are linked by direct transmission, *p*(*i* → *j*), we use a classification technique called naive Bayes. This method uses Bayes rule to estimate the probability of an outcome given a set of covariates from the observed frequencies in a training dataset. Our outcome, *L*_*ij*_ equals 1 if case *i* infected case *j* and 0 otherwise. We know the probable value of *L*_*ij*_ for case-pairs in the training set based on pathogen WGS or contact investigation data and want to predict the probability that *L*_*ij*_ = 1 for the remaining pairs.

We first use the training set to calculate *P*(*Z*_*k*_ = *z*_*k*_|*L* = *l*), the probability that the pair-level covariate *Z*_*k*_ equals *z*_*k*_ for each covariate *k* ∈ {1, 2, …, *p*} for a pair with link status *l* ∈ {1, 0}, using:

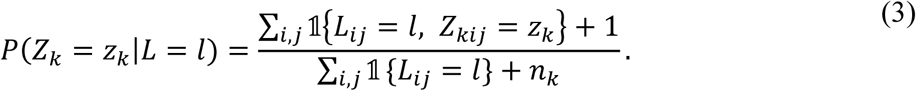

The indicator function, 𝟙, equals 1 if the input is true and 0 if false and *n*_*k*_ is the number of levels of *Z*_*k*_ for *k* ∈ {1, 2, …, *p*}. Therefore, the numerator, Σ_*i,j*_ 𝟙 {*L*_*ij*_ = *l, Z*_*kij*_ = *z*_*k*_}, counts how often a pair *i, j* has linked status *l* and value *z*_*k*_ for covariate *Z*_*k*_, *k* ∈ {1, 2, …, *p*}. Because sparse data can result in zero probabilities, we add 1 to each cell (33). Then we use the training set to calculate *P*(*L* = *l*), the prior probability of link status for *l* ∈ {1, 0} using:

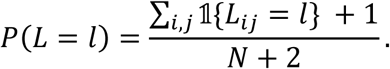

We use Bayes rule to calculate the predicted probability that case *i* infected case *j, p*(*i* → *j*) for all pairs in the prediction set as:

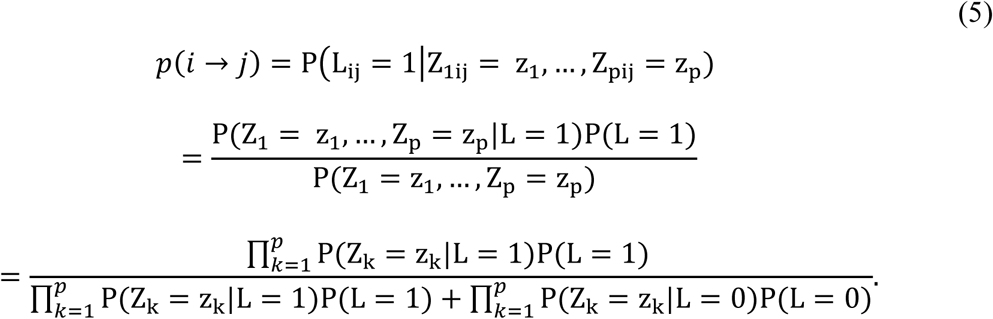

We calculate the conditional probability of all covariate values given link status P(Z_1_ = z_1_, …, Z_p_ = z_p_|L = 1), as the product of the conditional probabilities of each covariate, P(Z_k_ = z_k_|L = 1) for *k* ∈ {1, 2, …, *p*}, assuming that covariates are conditionally independent (i.e. that the covariate values are independent given link status).

Finally, the estimated probabilities are scaled to represent the relative likelihood that case *j* has been infected by case *i* rather than any other sampled case using:

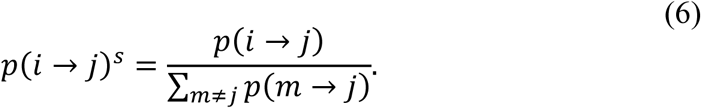

We call this scaled probability, *p*(*i* → *j*)^*s*^, the “relative transmission probability”. Note: the ordered nature of the pair dataset implies that if case *j* was observed before case *i*, then *p*(*i* → *j*) = 0.

### Training Dataset Construction

Naive Bayes uses a training set, with a known outcome, to inform a model to estimate probabilities in a separate prediction set. For many infectious diseases we never know the true infector. In our training set, the outcome therefore represents probable rather than certain transmission events, inferred from a subset of cases with pathogen WGS and/or detailed contact investigation data. Because of this uncertainty, we want to estimate the transmission probability of the training case-pairs as well as those which lack WGS or contact data and are only included in the prediction set. Therefore, we use an iterative estimation procedure where each pair has a turn in the prediction set. The algorithm used to create the training dataset and iterative estimation procedure is shown in Figure 1 and described further in the supplementary material.

**Figure 1.**
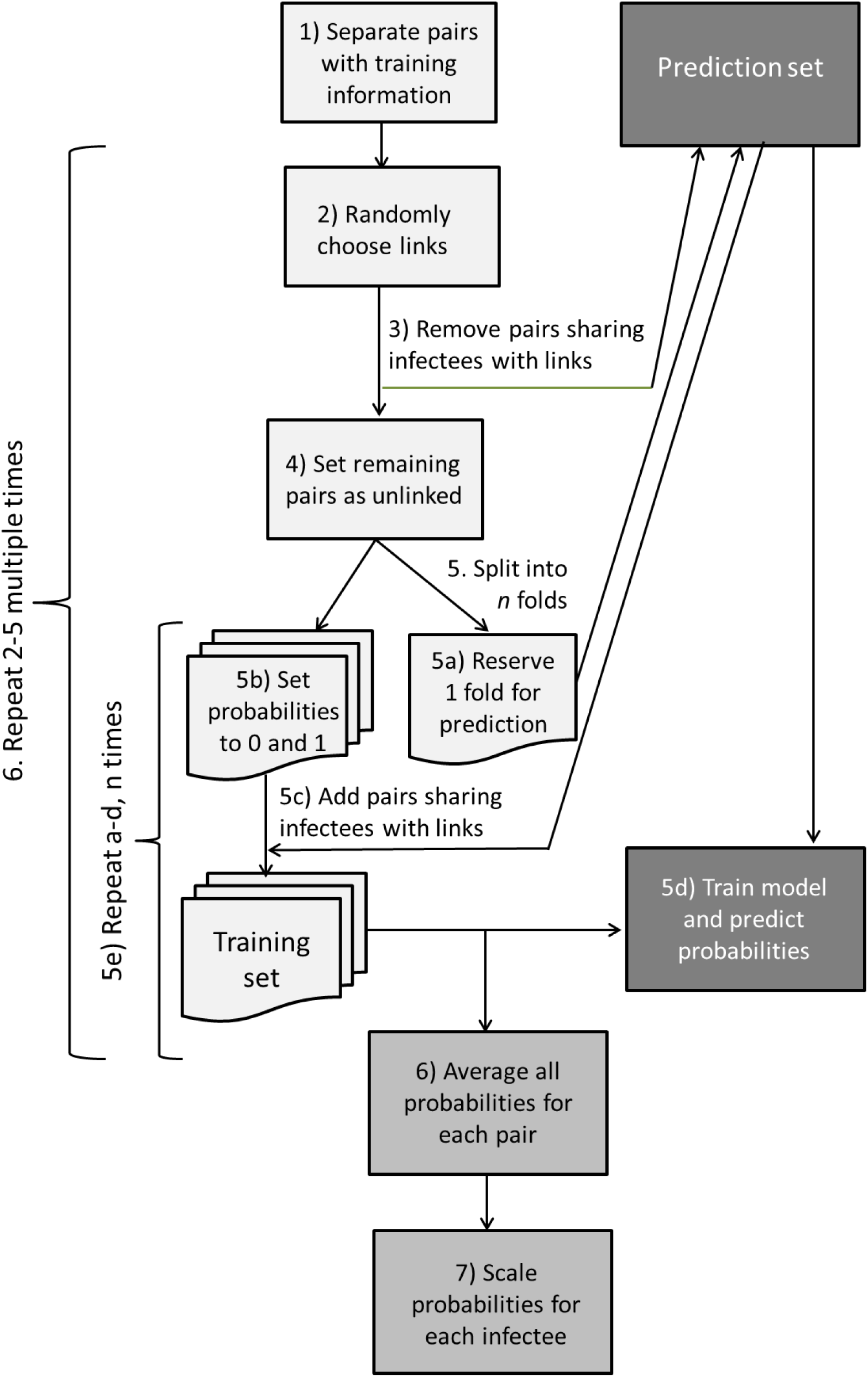
Flow-chart depicting the algorithm we used to create the training dataset and the iterative procedure to estimate the relative transmission probabilities.

### Reproductive Number Estimation

To estimate the reproductive number, we use the Wallinga and Teunis approach (4) that calculates the relative probability that each case was infected by all other cases using the serial interval distribution. The effective reproductive number (*R*_*i*_) for each case is then calculated by summing the scaled probabilities for all possible infectees assuming that all cases are sampled and the outbreak is completed with:

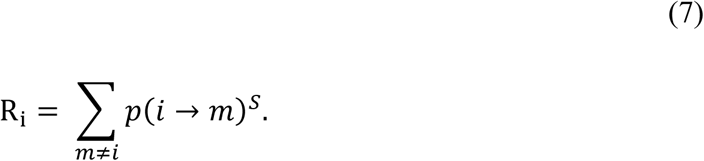

We use this equation, but with probabilities derived from our naive Bayes approach instead of probabilities based on the serial interval.

By averaging the individual reproductive numbers for all cases observed each month, we obtain monthly effective reproductive number (*R*_*t*_) estimates and average those values for the stable portion of the outbreak to give an average reproductive number estimate over the study period 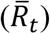. We calculate confidence intervals for *R*_*t*_ and 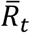 using parametric bootstrapping (see supplementary material).

### Simulation Study

We assess our method by applying it to 1000 simulated outbreaks of at least 500 cases, composed of multiple transmission chains using TB transmission dynamics (11,34). We simulate each transmission chain with an 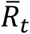 of 1.2 and a gamma distributed serial interval (shape=1.05, scale=2.0) (Ma, et al, under review). We simulate representative pathogen genomes and inform the model with four different covariates representing clinical and demographic variables and a discretized version of the time between infection.

We compare our method performance when training the model using probable transmission events defined by SNP distances, with training using truly linked and unlinked case-pairs. We also compare the method performance with that of probabilities derived from the time between infection dates and the serial interval distribution motivated by the Wallinga and Teunis method (4). For each simulation scenario (Table 1), we calculate the area under the receiver operating curve (AUC) and assess how the probability of the true infector rank compared to the probabilities of all possible infectors. We also estimate *R*_*t*_ and 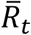. To determine what proportion of cases needs to be in the training set to achieve good performance, we use a sensitivity analysis with various outbreak sizes and training proportions. The simulation structure is explained further in the supplementary material.

### Hamburg TB Outbreak Application

We apply our method to a small TB outbreak in Hamburg and Schleswig-Holstein, Germany, analyzed in Roetzer et al. (10). The outbreak includes 86 individuals from the largest strain cluster in a long-term surveillance study conducted by the health departments in these cities. The dataset includes pathogen WGS data for all individuals as well as clinical, demographic, and social risk factor data. Furthermore, a subset of these individuals was involved in contact investigations performed by the local health authorities.

We define probable links in the training set in two ways: 1) SNP distances and 2) contact investigation. When training with SNP distance, case-pairs with <2 SNPs are considered linked, those with >12 SNPs are considered unlinked. Pairs with 2-12 SNPs are excluded from the training set as indeterminate (11,34). When using contact investigation data, pairs that had confirmed contact with each other are considered linked, pairs without confirmed contact with each other are considered unlinked, and cases who did not undergo contact investigation are excluded. For comparison, we also calculate the relative transmission probabilities randomly and using the same serial intervals as the simulation study.

## RESULTS

### Simulation Study

The sample sizes of the 1000 simulated outbreaks were 500-1178 (median: 545) and each outbreak had 2-39 (median: 14) individual transmission chains with 2-846 (median: 9) cases each. Supplementary Figure S1 shows the relative transmission probability distributions for one outbreak comparing truly linked and unlinked case-pairs. In that outbreak, our method estimated relative transmission probabilities of <0.005 for most unlinked pairs (92% when training with the truth and 89% training using SNP distance). With both ways of defining the training set, our method assigned more than 75% of truly linked case-pairs higher probabilities than the serial interval method (Supplementary Figure S2).

Over 1000 simulations, the average AUC was 97% (standard deviation [SD] 0.6) compared to 95% (SD 1.2) when the model was trained using true links and SNP distances respectively (Figure 2, Supplementary Table S1). When the model was trained with links determined by SNP distances, the estimated probability of the true infector ranked in the top 25% of all possible source case probabilities 93% (SD 2.4) of the time (compared to 95% (SD 1.6) when training with true links). Our method outperformed probabilities estimated using serial intervals (Figure 2, Supplementary Table S1). Figure 3 and Supplementary Table S2 show the 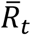 estimates for each of the different scenarios compared to the 1.2 value used to simulate the outbreaks. Both our method and the correct serial interval estimated 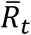 accurately. However, when incorrect serial intervals were used, the 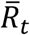 estimates were either too high or too low.

**Figure 2:**
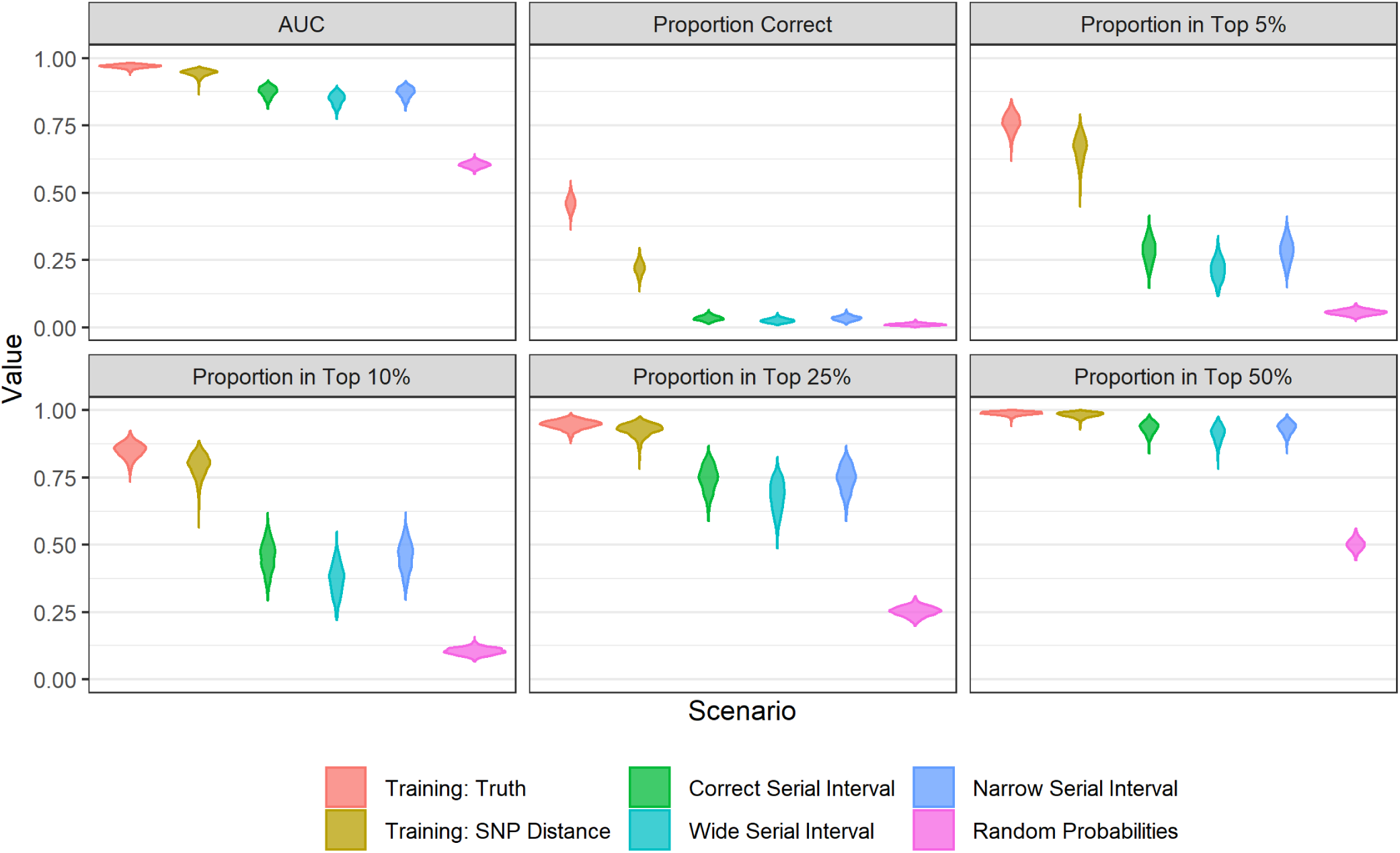
Violin plots of the performance metrics for the different scenarios across 1000 simulated outbreaks. The scenarios were: our method with a training set of true links, our method with a training set of links defined by SNP distance, probabilities derived from the serial interval distribution used to simulate the outbreak: gamma(1.05, 2.0), probabilities derived from a serial interval distribution that is too wide: gamma(1.3, 3.3) and too narrow: gamma(0.54, 1.9), and random probabilities. The metrics shown are the area under the receiver operating curve (AUC), the proportion of time the true infector was assigned the highest relative transmission probability (Proportion Correct), and the proportion of time the probability of the true infector was ranked in the top 5%, 10%, 25%, and 50% of all possible infectors.

**Figure 3:**
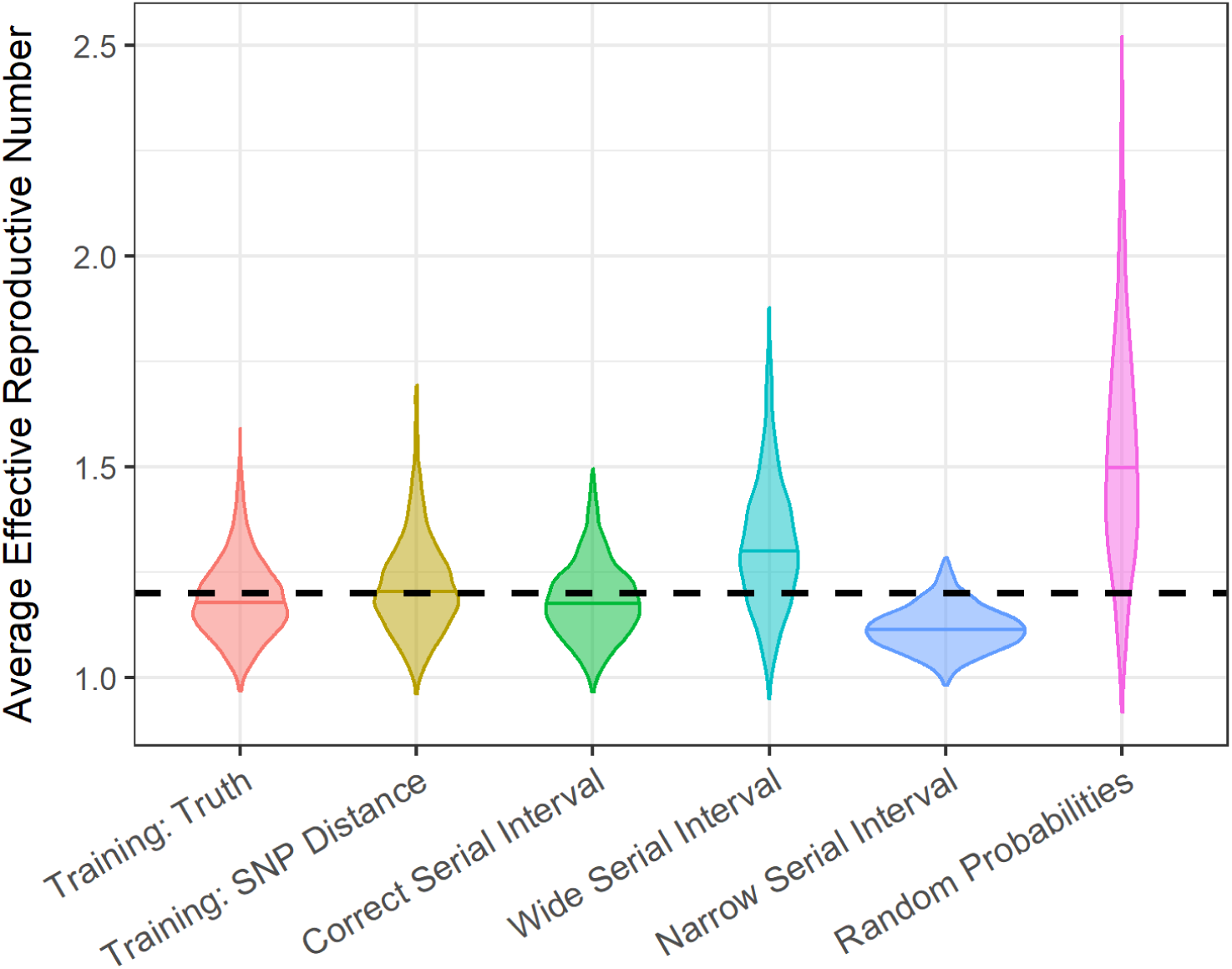
Violin plots of the distribution of the average effective reproductive number for different scenarios across 1000 simulated outbreaks. The dashed horizontal line indicates the true value of 1.2 that was used to simulate the outbreaks.

In our sensitivity analysis, the performance improved and the metrics’ variability decreased as the proportion of cases in the training set increased (Supplementary Figure S3). If the sample size was at least 500, only 10% of all cases were needed to train the model to obtain good performance. For a sample size of 200-500, training the model with 20% of cases resulted in good performance. For smaller outbreaks, the performance was best with at least 50% of the cases in the training set (Figure 4). The 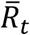 estimates grew increasingly accurate as the training dataset proportion increased and when 30% of cases were included, the estimates were close to the true value (Supplementary Figure S4).

**Figure 4:**
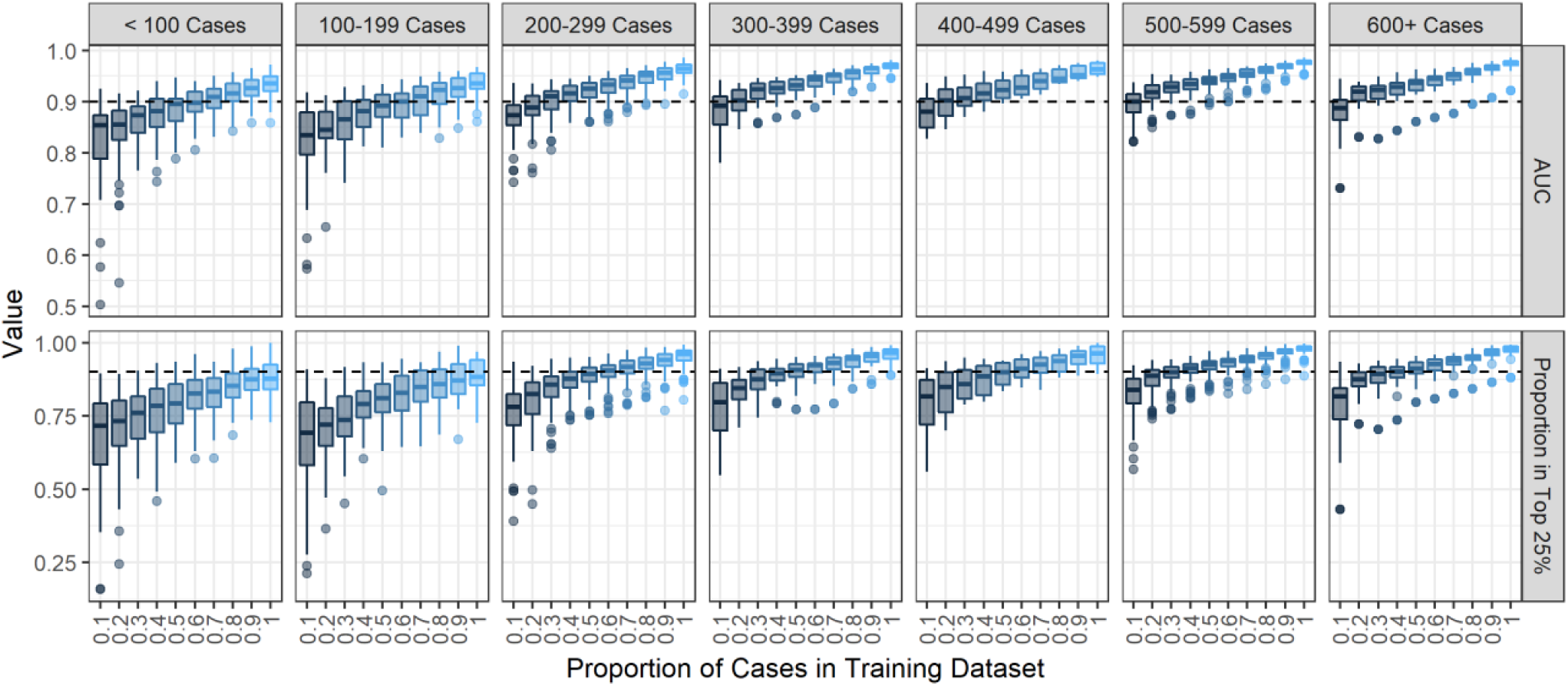
Boxplots of the performance metrics by training set proportion in 300 simulated outbreaks stratified by the total sample size of the outbreak. The metrics shown are the area under the receiver operating curve (AUC) and the proportion of time the relative transmission probability of the true source case was ranked in the top 25%. The dotted black line indicates a value of 90% on either metric.

### Hamburg TB Outbreak Application

Case counts over the course of the outbreak and clinical and demographic characteristics are shown in Figure 5 and Table 2. The 86 cases resulted in 3633 possible ordered case-pairs where the possible infector was observed before the infectee. These pairs were separated by 0-20 SNPs (median=4). Of the 86 individuals, 31 (36%) were part of contact investigations and 51 case-pairs had a confirmed contact. All individual-level covariates were transformed into pair-level covariates for analysis (Table 3).

**Figure 5:**
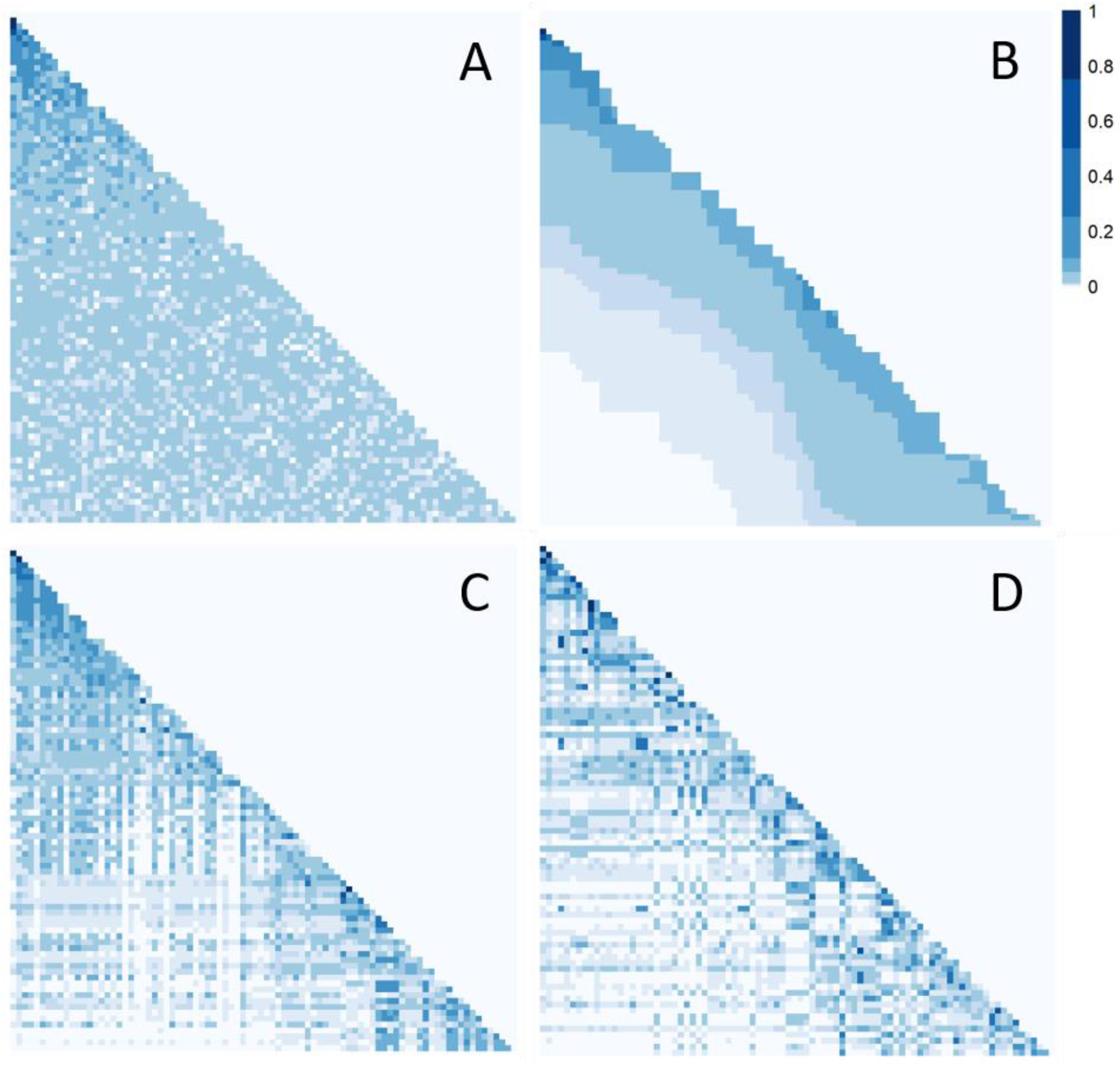
Case counts by year for the Hamburg outbreak described in Roetzer et al. (10)

Figure 6 shows heatmaps of all potential infectors for each infectee using our method compared to random probabilities (Figure 6A) and a serial interval distribution (Figure 6B). Using our method, defining links with either with SNP distance (Figure 6C) or confirmed contact (Figure 6D), there is more variation in the relative transmission probability across possible infectors than the serial interval or random scenarios. Some infectees have infectors with a higher probability than all others in the row, suggesting this is the likely true infector. However, even for rows without a clear single infector, many of the possible infectors have very low probabilities and can be eliminated as the true infector.

**Figure 6:**
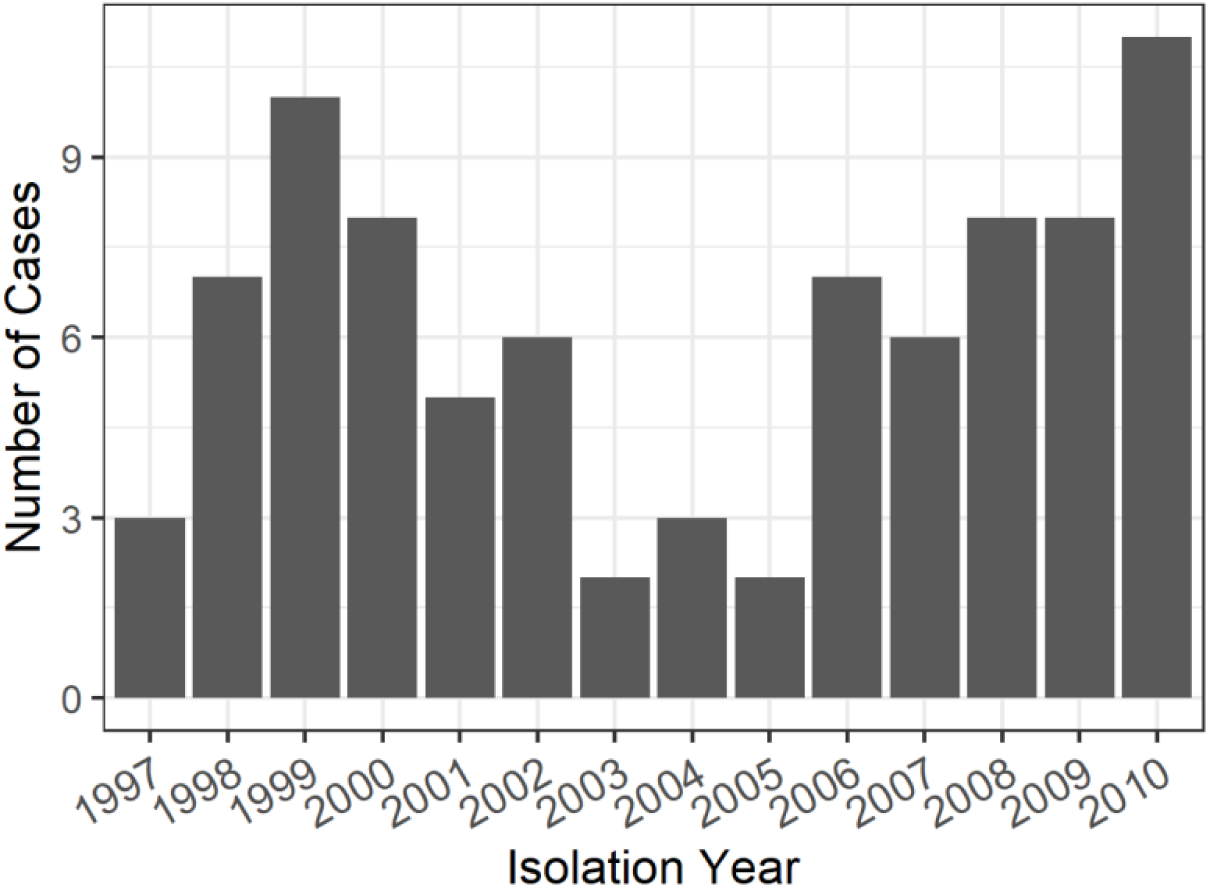
Heatmaps of the relative probabilities that each infectee (rows) was infected by each possible infector (columns) in the Hamburg TB outbreak. Darker squares represent higher the probabilities. The cases are ordered by infection date with the earliest cases on the top and to the left. Each panel shows the results from a different method of calculating probabilities: A) randomly assigned probabilities, B) probabilities calculated using a gamma(1.05, 2.0) serial interval distribution, C) probabilities calculated using our method and a training set with links based on SNP distance, and d) probabilities calculated using our method and a training set with links based on contact investigations.

All methods except random probabilities show spikes in *R*_*t*_ at the second peak in case counts, but to different degrees (Figure 7). The 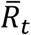 estimate was 0.97 (95% confidence interval [CI] 0.74-1.19) when training with contact investigation and 0.85 (95% CI 0.63-1.06) when training with SNP distances (Supplementary Table S3, Figure 8).

**Figure 7:**
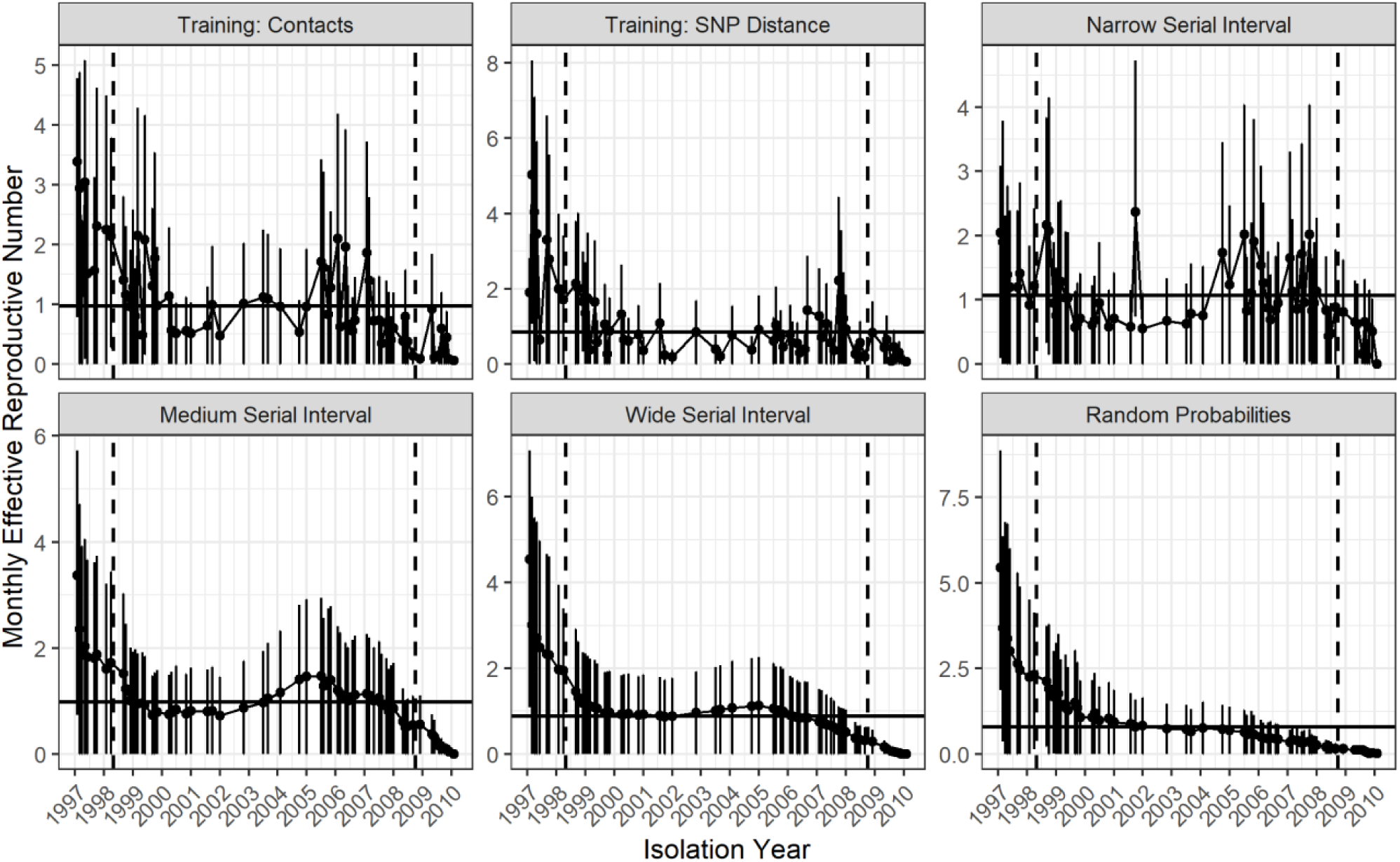
Monthly reproductive number over the course of the 14 years of the Hamburg TB outbreak estimated from the relative transmission probabilities with bootstrap confidence intervals. Each panel shows the results from a different method of calculating probabilities: our method and a training set with links based on contact investigation data; our method and a training set with links based on SNP distance; probabilities derived from narrow: gamma(0.54, 1.9), medium: gamma(1.05, 2.0), and wide: gamma(1.33, 3.0) serial interval distributions; and random probabilities. The months in between the dotted horizontal lines were averaged to find the average effective reproductive number for the scenario which is shown by the solid horizontal line.

**Figure 8:**
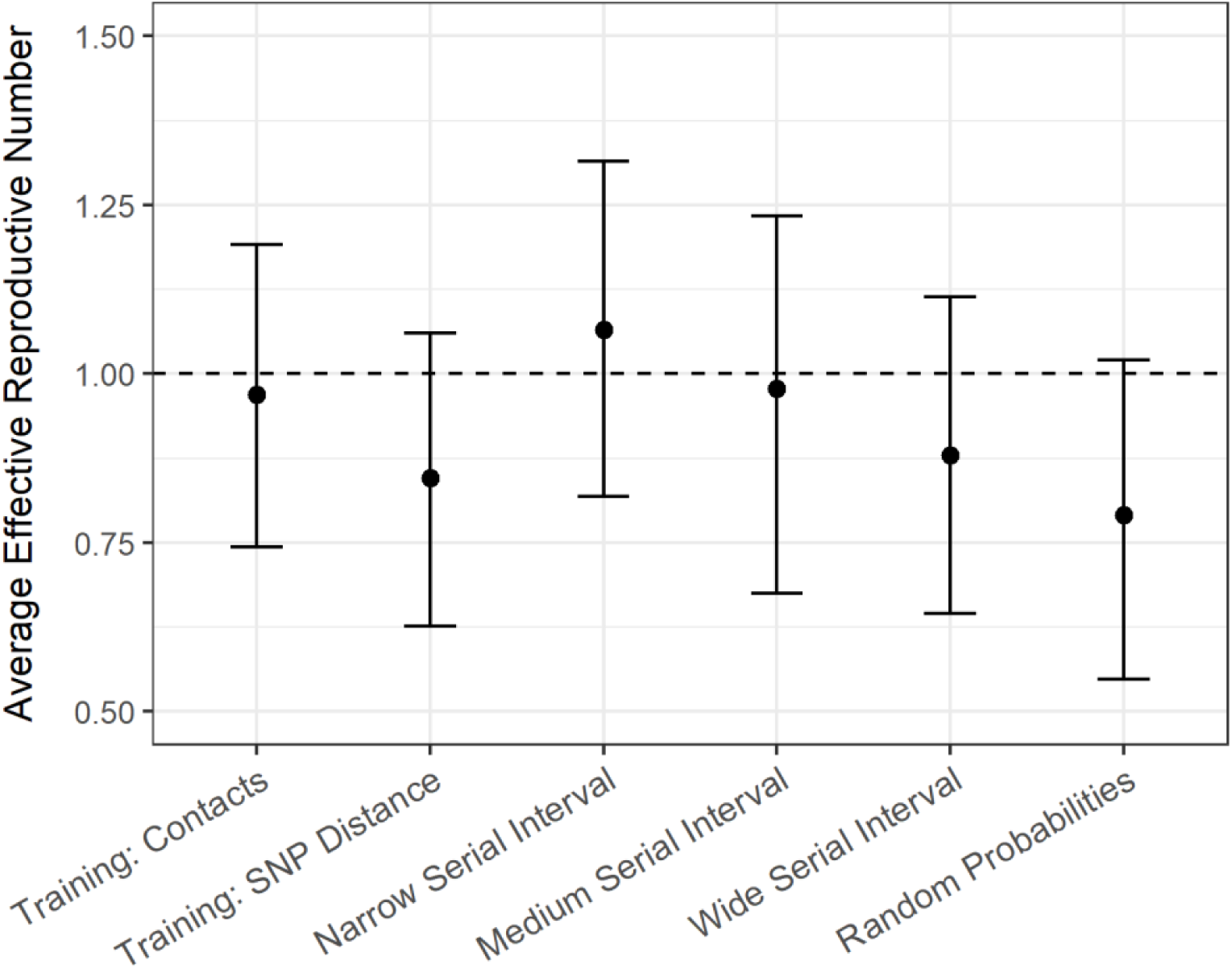
Average effective reproductive number for the Hamburg TB outbreak calculated using the relative transmission probabilities derived from different methods of calculating probabilities: our method and a training set with links based on contact investigation data; our method and a training set with links based on SNP distance; probabilities derived from narrow: gamma(0.54, 1.9), medium: gamma(1.05, 2.0), and wide: gamma(1.33, 3.0) serial interval distributions; and random probabilities. The vertical bars represent 95% bootstrap confidence intervals. The dotted horizontal line represents a 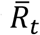 value of 1.

## DISCUSSION

We have developed a method to estimate the relative transmission probability between pairs of infectious disease cases using clinical, demographic, geographic, and genetic characteristics that accurately distinguishes between linked and unlinked case-pairs in simulation studies. Using a SNP distance proxy for transmission to train the model, the classification accuracy was 95%, and 93% of the time the true infector had a probability in the top 25% of all possible infectors. Our method outperformed the serial interval method in all metrics and accurately estimated 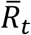. This is important because the serial interval is difficult to estimate and highly variable (7,8,35), highlighting the value of estimation methods that are independent of serial interval estimates.

Applying these methods to the Hamburg dataset, we found that both methods allowed for the elimination of many transmission links (Figure 6). The two ways of model training produced slightly different 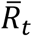 estimates, which is expected because neither of the different probable transmission events used to train the model perfectly capture the truth. Using contact investigation for training is more discriminating than SNP distance because we know the cases have actually interacted. However, this may miss links with unknown or unreported contacts. Using SNP distances for training will result in fewer missed links, but could connect cases that never had contact with one another. We hypothesize that the true reproductive number for *M. tuberculosis* in this context lies in between these two estimates (0.84-0.97).

Most established methods for exploring transmission focus on either identifying recent transmission clusters (11,16,34,36–39), recreating possible transmission chains (12,14,15,17–21,40–42), or identifying the true infector (43–47). When estimating transmission parameters, simply knowing clusters is not informative enough and identifying the true infector is often impossible in reality. The strength of our method is that it directly estimates the relative transmission probability for all case-pairs instead of seeking to find the true infector or a set of possible transmission trees. This gives our method broad applicability as it can identify potential true infectors (pairs with very high probabilities) or transmission clusters (groups of pairs with high probabilities). These probabilities can then be used to estimate transmission parameters incorporating the uncertainty around the true infector.

Other methods to estimate the transmission probabilities either only use genetic data (13) or require prior knowledge of the relationship between the covariates and transmission (48). Our method, however, uses many different information sources about the individual cases without assuming any relationship between these factors and transmission. Additionally, our method uses naive Bayes, a simple but powerful machine learning tool that has many diverse applications (33,49–52). Although traditionally a naive Bayes model is trained with a set of true events, our method performs almost as well when SNP distance is used as a transmission proxy.

Having both a training and prediction set also means that not all cases require highly discriminatory information such as contact investigation or pathogen WGS data to estimate relative transmission probabilities. This is relevant because existing datasets often have rich demographic, clinical, and spatial data but lack detailed contact investigation or pathogen WGS data due to significant time and resources needed to obtain these data. Provided a subset of cases, 10-50% depending on the sample size, has this information, our method can infer transmission patterns among the remaining cases as well.

Our method does make assumptions that might not be appropriate. Firstly, naive Bayes assumes independence of the covariates when conditioning on the outcome, which may not be realistic. However, numerous papers have shown that naive Bayes still performs well even when this assumption is violated (51,53–55). Furthermore, many naive Bayes extensions have been developed that relax this assumption (33,56,57), which could be easily integrated into our method.

The Wallinga and Teunis (4) approach for estimating 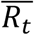 which we applied, assumes that every case was infected by someone that has been sampled. These authors and others found that simulations incorporating random incomplete reporting did not substantially decrease the accuracy of their 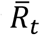 estimates, so this is unlikely to be an issue here (4,58). Our probability estimates themselves do not assume all cases in an outbreak are sampled because we estimate the relative probability that one case was infected by another over any other sampled case. However, our method could be affected by biased sampling, e.g. because only certain types of cases are observed or have the information needed to define probable links in the training set. Future work could more fully examine the effect of biased reporting and biased training sets.

Finally, as with other infectious disease analytical approaches, our method assumes that cases were infected in the same order that they were observed (39,59). Although not a strong assumption for diseases with clear symptoms and a short latent period, this may not be appropriate for diseases such as TB, with a highly variable and potentially long latent period, and often substantial delays in care-seeking care and diagnosis (60,61). Although this assumption is a known problem in infectious disease research, it is frequently made (43,44) because relaxing it complicates models substantially.

We have developed a method to estimate the relative transmission probabilities between pairs of cases, which is flexible, using any information sources available without making assumptions about the relationship between these covariates and transmission. The power of our method is that only a subset of cases requires pathogen WGS or contact investigation data, making this method applicable to many outbreak and surveillance datasets. These probabilities can be used to better understand the transmission dynamics of an outbreak by identifying or ruling out possible transmission events and estimating transmission parameters. In a disease where determining transmission events can be extremely difficult, using transmission probabilities between all possible cases provides a unique and powerful analysis tool.

## FUNDING

This work was supported by the US National Institutes of Health [NIHGMS R01GM122876]. SVL was also funded by the Interdisciplinary Training grant from the US National Institutes of Health [NIHGMS T32GM074905]. HEJ was also funded by US National Institutes of Health [NIH K01AI102944]. CRH and LFW were also funded by the Providence/Boston Center for AIDS Research [P30AI042853]. CRH was supported by the Boston University/Rutgers Tuberculosis Research Unit [U19AI111276] and the U.S.-India Vaccine Action Program (VAP) Initiative on Tuberculosis (CRDF Global/NIAID). RSL holds a Fellowship from the Canadian Institutes of Health Research [MFE-152448]. The content of the article is solely the responsibility of the authors and does not necessarily represent the views of the National Institute of Allergy and Infectious Disease or the Office of the Director, NIH. The funders had no role in the decision to publish this manuscript.

## Notes

#### Summary of Updates

Paper adapted to journal submission and algorithm updates changed results slightly.

