## supplementary material for "Estimating the Relative Probability of Direct Transmission between Infectious Disease Patients"

### NAIVE BAYES DETAILS

#### *Covariate Distances*

When we transform our starting dataset of individuals into a dataset of ordered pairs, we convert the individual-level covariates ($\mathrm{X}_1, \mathrm{X}_2, \dots, \mathrm{X}_\mathrm{p}$) into pair-level covariates ($Z_1, Z_2, \dots, Z_p$) by computing the "distances" between the covariate values for the two cases. These distances can be dichotomous where a value of 1 indicates that the covariate values match and 0 indicates that they do not match as in Equation 1 for covariate $Z_k$ (where $k \in \ 1, 2, \dots, p$):

$$Z_{kij} = d(X_{ki}, X_{kj}) = \begin{cases} 1 \ if \ X_{ki} = X_{kj} \\ 0 \ if \ X_{kj} \neq X_{kj} \end{cases} . \quad (1)$$

Or the distances could be categorical where different combinations of individual-level values result in $n_k$ different pair-level values such as:

$$Z_{kij} = d(X_{ki}, X_{kj}) = \begin{cases} z_{k1} \ if \ X_{ki} = X_{kj} = x_{k1} \\ z_{k2} \ if \ X_{ki} = X_{kj} = x_{k2} \\ z_{k3} \ if \ X_{ki} = x_{k1}; \ X_{kj} = x_{k2} \\ \dots \\ z_{kn_k} \ if \ X_{ki} = X_{kj} = x_{kw} \end{cases} \quad (2)$$

where $x_{k1}, \ x_{k2}, \dots, \ x_{kw}$ are the different values of $X_k$ and $z_{k1}, \ z_{k2}, \dots, z_{kn_k}$ are the different values of $Z_k$ .

#### *Training Dataset Creation Algorithm*

To estimate the transmission probabilities, we split our training set into n subsets called folds where n - 1 folds are used to train the model and the remaining fold is included in the prediction set. The fact that we assume that each case has only one true infector presents two problems for the creation of a proper training set: 1) since the method of defining transmission events in the training set is not perfect, there could be multiple possible links for each infectee, and 2) once we denote a case pair as linked in the training set, all other possible pairs that share that infectee have a zero-probability of being linked.

To solve the first problem, we create the links in the training set by randomly choosing one of the possible infectors defined by pathogen WGS or contact investigation data to be designated the linked case-pair for this run and then repeat this selection multiple times to capture the uncertainty around the true infector. To solve the second problem, when a pair is designated as a link in the training set, all other appropriately timed pairs with the same infectee as the link are also included in the training set as non-links. This procedure means that when a linked pair is in the prediction dataset so are all of the other pairs involving the infectee, so the probability of all pairs possible involving that infectee can be estimated.

In order to capture the valuable information provided by the pathogen WGS or contact investigation data used to define probable transmission events in the training set, when a pair is denoted as linked or unlinked for a particular run by that information, the predicted probability for that run is set to 1 if the pair is linked and 0 if the pair is not linked. Note the probability is not set to 0 for those pairs that are just included in the training dataset because they share an infectee with a designated link. The following algorithm describes the training dataset creation and iterative estimation procedure:

1. Create a dataset of possible training case pairs by subsetting the dataset to only the pairs involving individuals that have information used to define probable links in the training set.

2. Randomly select one infector for each infectee out of all possible infectors and designate those pairs as “linked” ($L_{ij} = 1$).
3. Temporarily remove all pairs that share an infectee with the linked pairs defined in step 2 from the training set.
4. Designate all remaining unlinked pairs as not linked ($L_{ij} = 0$).
5. Split this dataset of possible training pairs into n folds (we used 10): n-1 for training and 1 for prediction.
    a. Reserve 1 fold for prediction and combine with all of the pairs not in the training set.
    b. Set the predicted probabilities for training set pairs to 1 for links and 0 for non-links.
    c. For all linked pairs in the n-1 training folds, move all other pairs involving the infectee from the prediction set to the training set as unlinked ($L_{ij} = 0$). This is the final training set for this iteration.
    d. Use the training set to train the model and calculate predicted probabilities in the prediction set.
    e. Repeat (a)-(d) n times so that each fold has a turn in the prediction set.
6. Repeat steps 2-5 multiple times (we used 10) to allow for different possible infectors to be designated the true infector.
7. Average over all the predicted probabilities for each pair.
8. Scale the resulting probabilities using equation 6 to obtain the relative transmission probabilities for all case pairs.

*Bootstrap confidence intervals for the reproductive number*

In order to estimate confidence intervals for $R_t$ and $\overline{R_t}$ we use parametric bootstrapping. First, we re-sample the $R_i$ values 100 times using the estimated probabilities, $p(i \rightarrow m)^S$, for all possible infectees according to their distribution:

$$R_i \sim \sum_{m \neq i} Bernoulli(p(i \rightarrow m)^S) \quad (1)$$

as detailed in (1). For each re-sampling we calculate the $R_t$ values at each month and average them to estimate $\overline{R_t}$. We then use the distributions of the estimated values of $R_t$ and $\overline{R_t}$ to derive 95% bootstrap confidence intervals:

$$\text{LowerBound} = \hat{\text{R}} - (\text{Q}_{\tilde{R}}(1 - \alpha/2) - \hat{\text{R}}) \quad (2)$$

$$\text{UpperBound} = \hat{\text{R}} - (\text{Q}_{\tilde{R}}(\alpha/2) - \hat{\text{R}})$$

where $\hat{R}$ can be either$\widehat{\overline{R_t}}$ or $\widehat{R_t}$ and $\text{Q}_{\tilde{R}}$ is the quantile function of the bootstrap estimates of $R$.

SIMULATION DETAILS

*Initial set-up*

We simulate 1000 outbreaks using the **simulateOutbreak** function and generate the phylogenetic trees for those outbreaks using the **phlyoFromPTree**, function both from *TransPhylo* v1.0 (2). Then we use the **simSeq** function in *phagnorn* v2.5.3 (3) to generate genetic sequences corresponding to the phylogenetic tree. This method was used in Stimson et al. (4) for a similar purpose, though we extend it to include multiple transmission chains. Each chain has at least two cases and we simulate the chains iteratively until the total sample size exceeds

500. The different transmission chains all last for 20 years but have start and end points that vary in time. We used an effective population size times generation time ($N_{eg}$) of 0.25.

Although pathogen genomes are thousands to millions of base-pairs long, for computational efficiency, we simulate a 300 base pair genome where each transmission chain starts with a unique set of base pairs to allow for genetic diversity across different strains. The shortened genome length we simulate is meant to represent the locations that differ amongst cases sampled as part of one outbreak instead of the full genome. This simplification is appropriate because we aim to replicate a slow mutating pathogen such as TB which mutates at a rate of around 0.5 SNPs per genome per year (5). With this mutation rate, over the course of one 20 year transmission chain, very few mutations will accrue thus allowing a smaller genome to provide a good proxy for the full genome. We also performed a sensitivity analysis which showed that the SNP distance for a fixed mutation rate did not notably change as genome length increased (Figure S5). It is possible that this approach underestimates the true SNP distance distribution which makes it a conservative representation of the true SNP distance distribution using the full pathogen genome.

*Covariate Simulation*

We added covariates to the outbreak structure produced by **TransPhylo** that were associated with link status. We simulated four covariates at the individual level, $X_1 - X_4$, with pair level analogs, $Z_1 - Z_4$. The covariate values for each source case were chosen randomly using the population frequencies (Table S1). We looped through the pairs starting with the earliest case with a sampled infector and chose the covariate values for that case based on the values for their infector according to the rules set for that particular covariate (Table S1). We also included the time between infection dates for each case pair into the model categorized as follows: less than 1 year, 1 to 2 years, 2 to 3 years, 3 to 4 years, 4 to 5 years, and more than 5 years.

These covariates were chosen arbitrarily simply to create covariates with different structures that had different magnitudes of association with whether or not a pair was linked. However, $X_1/Z_1$ could represent the presence of some social risk factor where we believe that pairs who both have or do not have the risk factor are more likely to be linked. Covariate, $X_2/Z_2$ could represent nationality where there are multiple groups and we believe that pairs with the same nationality are more likely to be linked. Covariate, $X_3/Z_3$ could represent a age group where we believe that the characteristic of the infector matters, so order is important. Finally, $X_4/Z_4$ could represent geographic location where pairs that live in the same or close areas are more likely to be linked.

*Simulation Schemes*

We estimated the relative transmission probabilities using six different scenarios: naive Bayes training with true links and training with links derived from SNP distance, three different serial interval distributions (correct, wide and narrow), and random assignment. When we train the model using SNP distance, case pairs with less than 2 SNPs are considered linked and those with more than 12 SNPs are considered unlinked in the training set (11,34). Pairs with between 2 and 12 SNPs are considered indeterminate and thus are not included in the training set. To represent real scenarios when only a fraction of the cases have the discriminating information necessary to define probable transmission events, we randomly select a subset of 60% of all cases to make up the training set both when training with true links and links derived from SNP distances.

The serial interval comparison was motivated by the Wallinga and Teunis (4) method for calculating the reproductive number in which the transmission probabilities are estimated from the serial interval distribution. The wide and narrow serial intervals represent the prior and posterior distributions used by Didelot et al. (35) which were chosen because they were derived from the same TB outbreak in Hamburg that we analyze below. As a negative control, we assign the probabilities randomly from a Uniform(0, 1) distribution. Note that because in real datasets, we do not know the infection dates, but in simulations we do, when we refer to the "serial interval" we are referring to the generation interval in the simulation studies and the serial interval in real applications. Additionally all of these serial intervals are shifted by 90 days because a serial interval of less than 3 months is not possible for TB, our motivating example.

We assess how well the relative transmission probabilities could classify case pairs as linked and unlinked across all 1000 simulated outbreaks, by calculating the area under the receiver operating curve (AUC) for each simulation. Additionally to determine how well our method performed in identifying the true infector, we evaluated how the relative transmission probability of the true infector ranked compared to the probabilities of all possible infectors. We also perform a sensitivity analysis to determine what proportion of cases needed to be included in the training set to achieve good performance. We simulate 300 outbreaks with sample sizes ranging from 50-1000 cases and for each outbreak we perform our method assuming that a random subset of 10% to 100% of the cases can be included in the training set. We assess how changing the proportion of cases sampled affects the performance of the model performance and the estimation of the reproductive number depending on the sample size.

Table S1. Covariate Simulation Scheme

| Variable | Frequency | Paired Version | Linked Pairs |
|---|---|---|---|
| $X_1/Z_1$ | a: 50% b: 50% | $Z_1$ = 1 if same<br>$Z_1$ = 0 if different | 60% chance of same<br>40% chance of different |
| $X_2/Z_2$ | a: 50% b: 30%<br>c: 15% d: 5% | $Z_2$ = 1 if same<br>$Z_2$ = 0 if different | 70% chance of same<br>30% chance of different |
| $X_3/Z_3$ | a: 70% b: 30% | $Z_3$ = 1 if a-a<br>$Z_3$ = 2 if b-b<br>$Z_3$ = 3 if a-b<br>$Z_3$ = 4 if b-a | Infector is a: 80% a-a<br>20% a-b<br>Infector is b: 70% b-b<br>30% b-a |
| $X_4/Z_4$ | a: 5% b: 5%<br>c: 5% d: 20%<br>e: 30% f: 10%<br>g: 10% h: 5%<br>i: 5% j: 5% | $Z_4$ = 1 if same<br>$Z_4$ = 2 if neighbors<br>$Z_4$ = 3 if other | 60% chance of same<br>35% chance of neighbors<br>5% chance of other |

Table S2. Performance metrics for relative transmission probabilities over 1000 simulations

| Scenario | AUC<br>Percent (SD) | Correct[1]<br>Percent (SD) | Top 5%[2]<br>Percent (SD) | Top 10%[2]<br>Percent (SD) | Top 25%[2]<br>Percent (SD) | Top 50%[2]<br>Percent (SD) |
|---|---|---|---|---|---|---|
| Gold Std: Truth | 96.9 (0.6) | 46.1 (2.6) | 76.1 (3.5) | 85.1 (2.9) | 94.7 (1.6) | 98.8 (0.7) |
| Gold Std: SNP Distance | 94.6 (1.2) | 22.1 (2.5) | 66.4 (5.0) | 79.5 (4.3) | 92.7 (2.4) | 98.3 (0.9) |
| Correct Serial Interval | 87.5 (1.9) | 3.4 (0.9) | 28.5 (4.4) | 45.6 (5.5) | 74.6 (4.9) | 93.5 (2.2) |
| Wide Serial Interval | 84.8 (2.2) | 2.5 (0.7) | 21.5 (4.0) | 37.1 (5.7) | 68.1 (5.9) | 91.4 (2.9) |
| Narrow Serial Interval | 87.3 (1.9) | 3.4 (0.9) | 28.5 (4.4) | 45.6 (5.5) | 74.6 (4.9) | 93.5 (2.2) |
| Random Probabilities | 60.4 (1.1) | 1.0 (0.4) | 5.6 (1.0) | 10.5 (1.3) | 25.4 (1.8) | 50.2 (2.0) |

[1]Percentage of the time the true infector for each case is assigned the highest probability of all possible infectors

[2]Percentage of the time the probability the probability of the true infector for each case is in the top 5, 10, 25, or 50% of all possible infectors

Table S3. Average effective reproductive number for different simulation scenarios

| Scenario | $\overline{R_t}$, mean (SD) |
|---|---|
| Training: Truth | 1.19 (0.09) |
| Training: SNP Distance | 1.22 (0.11) |
| Correct Serial Interval | 1.18 (0.09) |
| Wide Serial Interval | 1.31 (0.15) |
| Narrow Serial Interval | 1.12 (0.05) |
| Random Probabilities | 1.53 (0.27) |

Table S4. Average effective reproductive number for Hamburg outbreak by method

| Scenario | $\overline{R_t}$, mean (95% CI) |
|---|---|
| Gold Std: Confirmed Contact | 0.97 (0.74, 1.19) |
| Gold Std: SNP Distance | 0.85 (0.63, 1.06) |
| Narrow Serial Interval | 1.07 (0.82, 1.31) |
| Medium Serial Interval | 0.98 (0.68, 1.23) |
| Wide Serial Interval | 0.88 (0.64, 1.11) |
| Random Probabilities | 0.79 (0.55, 1.02) |

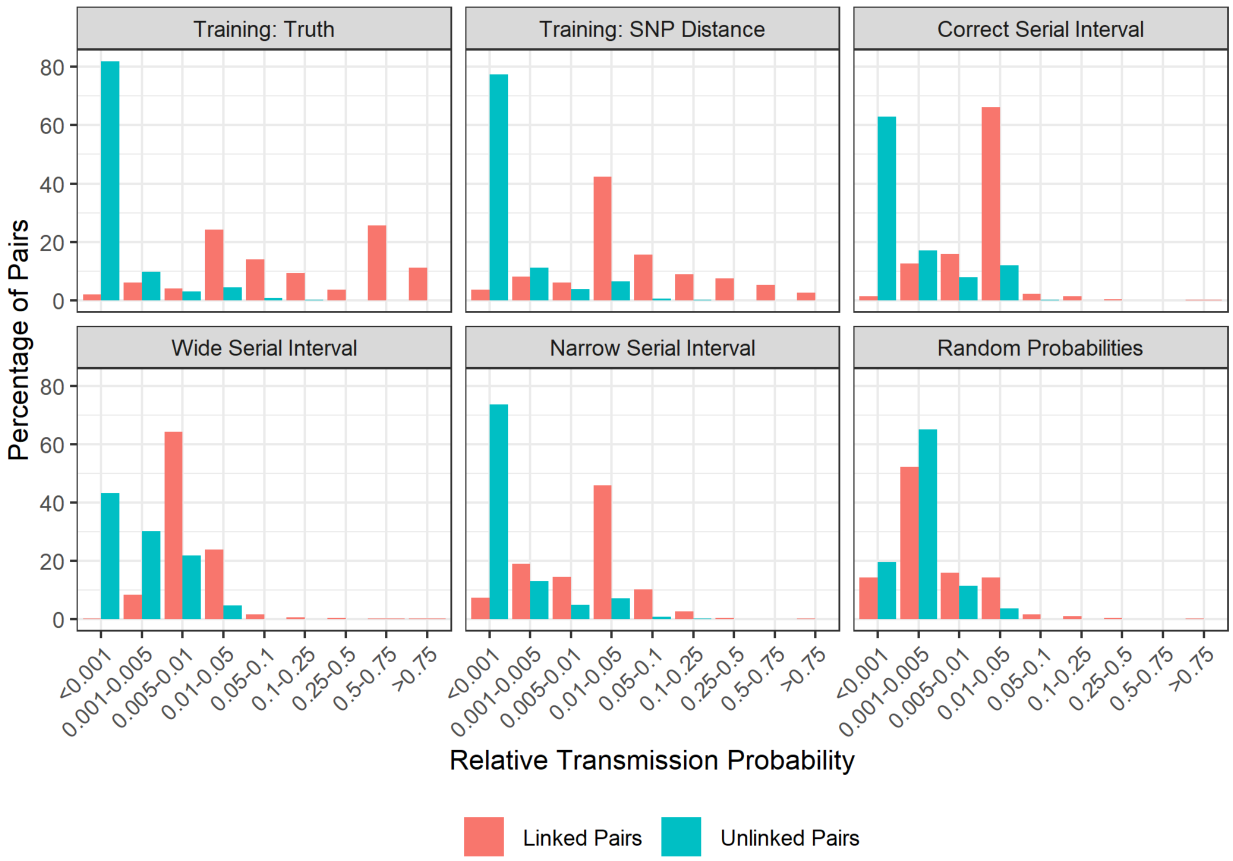


Figure S1. Distribution of the relative transmission probability – how much more likely is it that case $i$ was infected by case $j$ as opposed to any other sampled case – for linked (black) and unlinked (grey) case pairs in one of the 1000 simulated outbreaks. Each panel shows a different method of calculating probabilities: our method with a training set of true links, our method with a training set of links defined by SNP distance, probabilities derived from the serial interval distribution used to simulate the outbreak: gamma(1.05, 2.0), probabilities derived from a serial interval distribution that is too wide: gamma(1.3, 3.3) and too narrow: gamma(0.54, 1.9), and random probabilities.

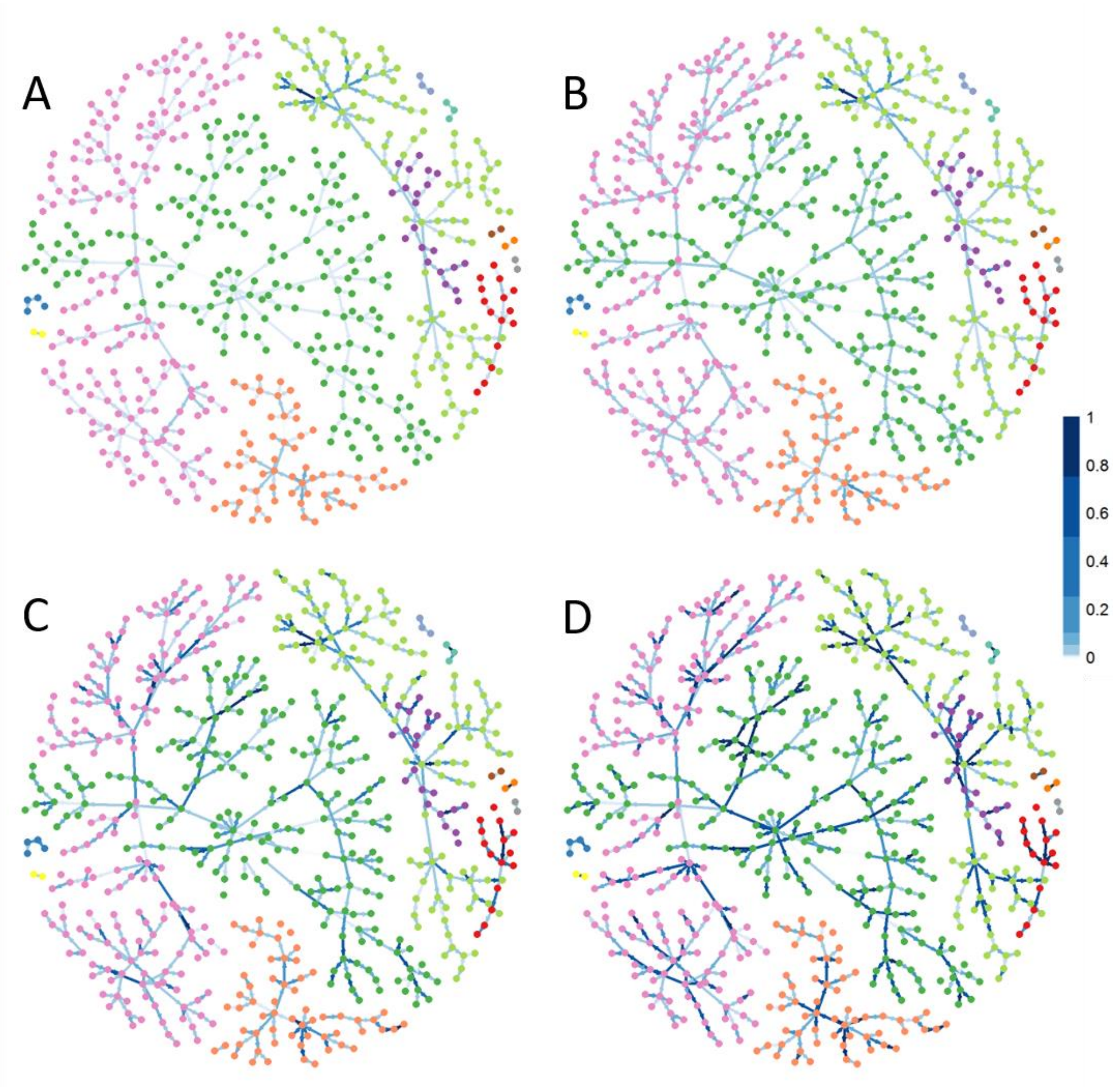


Figure S2. Network plots of the true transmission network in one of the 1000 simulated outbreaks. The nodes represent individual cases and are colored by transmission chain. The edges represent true transmission events and are colored based on the estimated relative transmission probability; the darker the color the higher the probabilities. A) Edges colored based on randomly assigned probabilities. B) Edges colored based on probabilities calculated by the correct serial interval: gamma(1.05, 2.0). C) Edges colored based on the probabilities calculated using our with a training set of links defined by SNP distance. D) Edges colored based on the probabilities calculated using our method with a training set of true links.

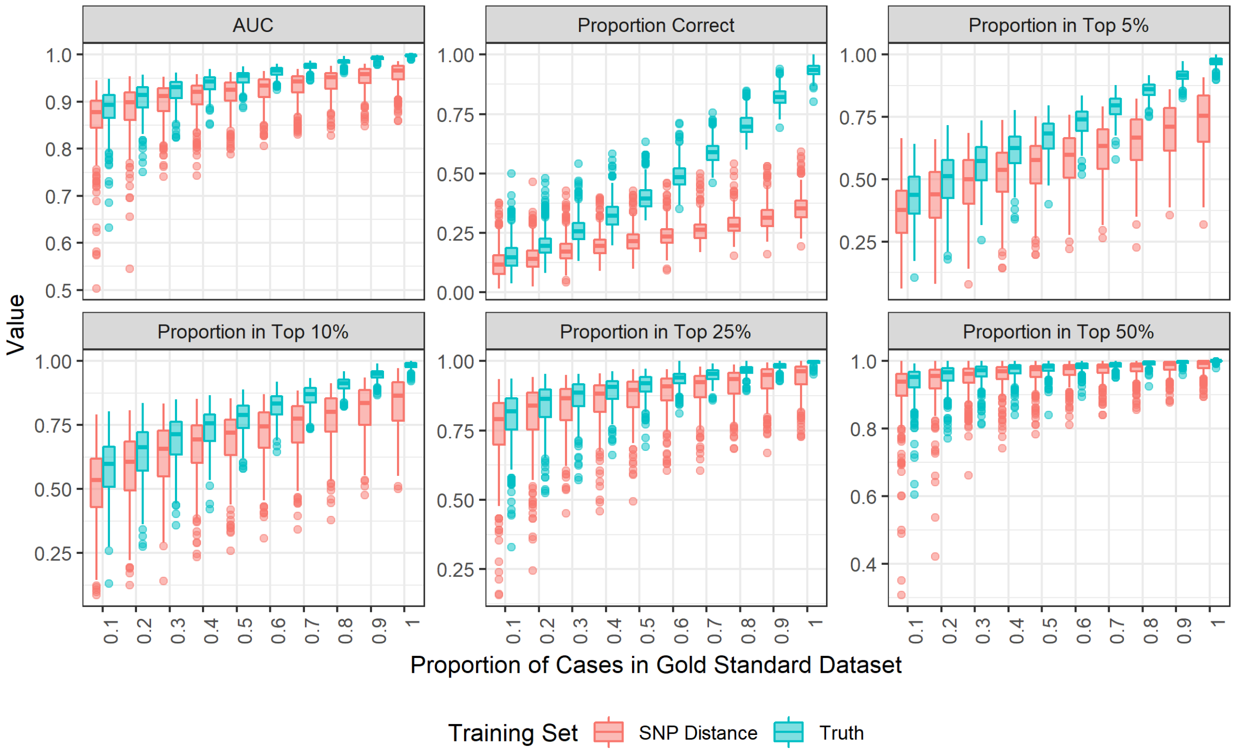


Figure S3. Boxplots of the performance metrics when varying the training set proportion in 300 simulated outbreaks with sample sizes varying from 50-1000. The plots are colored by the type of gold standard: SNP distance (red) or true transmission (blue). The metrics shown are the area under the receiver operating curve (AUC), the proportion of time the true infector was assigned the highest relative transmission probability (Proportion Correct), and the proportion of time the probability of the true infector was ranked in the top 5\%, 10\%, 25\%, and 50\% of all possible infectors.

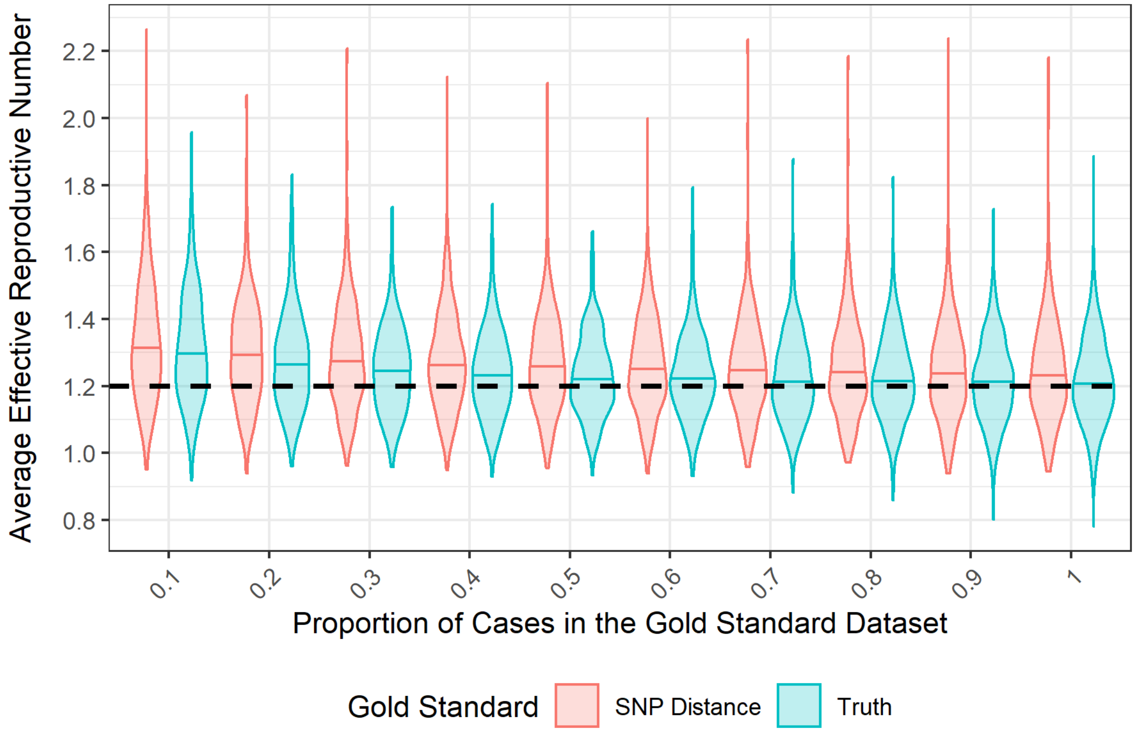


Figure S4. Violin plots of the distribution of the average effective reproductive number when varying the dataset proportion in 300 simulated outbreaks with sample sizes varying from 50-1000. The plots are colored by the way of defining links in the training set: SNP distance (red) or true transmission (blue). The dashed horizontal line indicates the true value of 1.2 that was used to simulate the outbreaks.

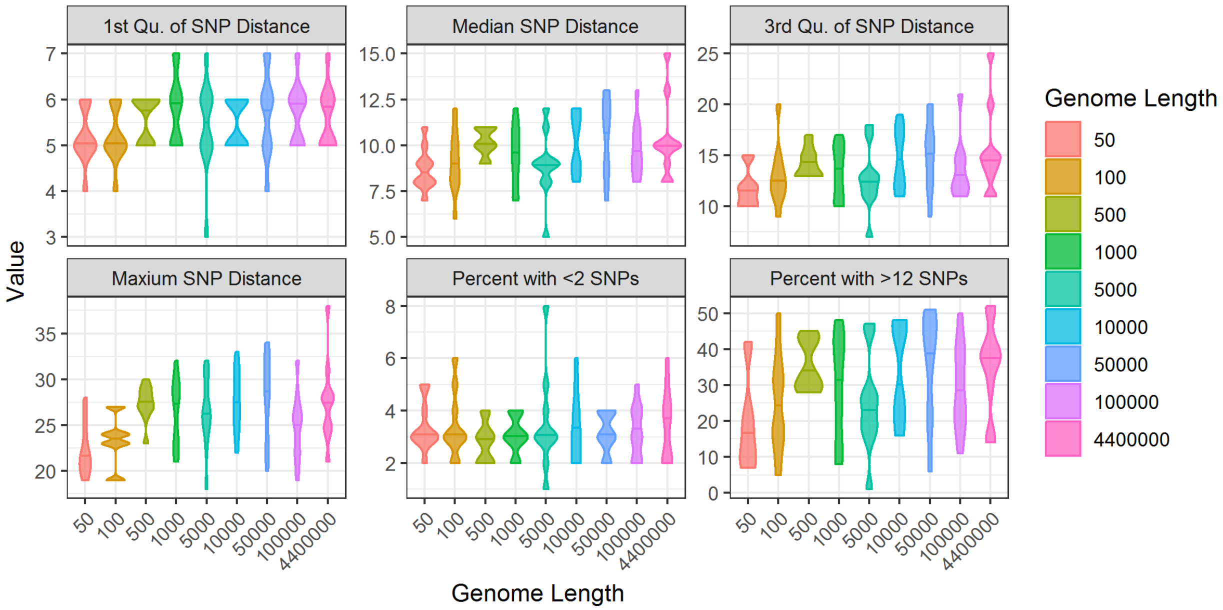


Figure S5. Violin plots representing the effect of changing genome length on the SNP distance distribution between case pairs. Pathogen genomes of various lengths from 50 to 4.4 million base pairs were simulated 100 times each for an outbreak of 200 cases. The resulting SNP distance matrix for all case pairs was computed. The figure shows the relationship between genome length on the quartiles of the SNP distance distribution as well as the maximum SNP distance and the percent of pairs with less than 2 SNPs and more than 12 SNPs.

REFERNCES
